# The mutational signatures of formalin fixation on the human genome

**DOI:** 10.1101/2021.03.11.434918

**Authors:** Qingli Guo, Eszter Lakatos, Ibrahim Al Bakir, Kit Curtius, Trevor A. Graham, Ville Mustonen

**Affiliations:** Organismal and Evolutionary Biology Research Programme, Department of Computer Science, University of Helsinki, 00014 Helsinki, Finland; Evolution and Cancer Laboratory, Centre for Genomics and Computational Biology, Barts Cancer Institute, Barts and the London School of Medicine and Dentistry, Queen Mary University of London, Charterhouse Sq, London, EC1M 6BQ, UK; Institute of Biotechnology, Helsinki Institute for Information Technology, University of Helsinki, 00014 Helsinki, Finland

**Keywords:** Formalin fixation paraffin embedding (FFPE), mutational signatures, artefact correction, computational genomics

## Abstract

**Background:** Formalin fixation and paraffin embedding (FFPE) of patient material remains standard practice in clinical pathology labs around the world. Clinical archives of patient material near-exclusively consist of FFPE blocks. The ability to perform high quality genome sequencing on FFPE-derived DNA would accelerate a broad spectrum of medical research. However, formalin is a recognised mutagen and sequencing of DNA derived from FFPE material is known to be riddled with artefactual mutations.

**Results:** Here we derive genome-wide mutational signatures caused by formalin fixation, and provide a computational method to correct mutational profiles for these formalin-induced artefacts. We show that the FFPE-signature is dominated by C>T transitions caused by cytosine deamination, and has very high similarity to COSMIC signature SBS30 (base excision repair deficiency due to inactivation mutations in *NTHL1*). Further, we demonstrate that chemical repair of formalin-induced DNA lesions, a process that is routinely performed as part of sequencing library preparation, leads to a signature highly similar to COSMIC signature SBS1 (spontaneous deamination of methylated cytosine). Next, we design FFPEsig, a computational method to remove the formalin-induced artefacts from mutational counts. We prove the efficacy of this method by generating synthetic FFPE samples using 2,780 cancer genomes from the Pan-Cancer Analysis of Whole Genome (PCAWG) project, and via analysis of FFPE-derived genome sequencing data from colorectal cancers.

**Conclusions:** Formalin fixation leaves a predictable mutational footprint across the genome. The application of our FFPEsig software corrects the mutational profile for the influence of formalin, enabling robust mutational signature analysis in FFPE-derived patient material.

## Background

Patient samples are routinely processed with formalin fixation and paraffin embedding (FFPE) by pathology laboratories around the world. FFPE preserves tissue morphology and enables immunohistochemical analysis for clinical diagnosis [1,2]. However, genomic analysis of DNA extracted from FFPE blocks is problematic, as formalin fixation negatively impacts DNA quantity and quality compared to fresh frozen (FF) material [3,4]. The pathology archive of any large hospital is likely to contain tens of thousands of FFPE blocks. Enabling accurate genomic analysis of FFPE material would unlock the tremendous translational research potential of these vast collections of archival material.

During fixation step of FFPE preservation, buffered formalin (4% formaldehyde) penetrates the biospecimen and generates cross-links between intracellular macromolecules (DNA-DNA, DNA-RNA and DNA-protein). These crosslinks stall DNA polymerases during library amplification [5–7]. As a consequence, the diversity and the number of templates that can be amplified by PCR from FFPE DNA is significantly depleted [4,8]. Furthermore, formalin causes hydrolytic deamination of cytosine bases to uracil [1,5], resulting in U:G mismatches where DNA polymerase incorporates adenine opposite to uracil in amplicon-based protocols, generating artefactual C:G>T:A substitutions in sequencing data [5,9,10].

To mitigate deamination artefacts, some FFPE sequencing library preparations include “repair treatment” whereby uracil DNA glycosylase (UDG) is added to remove uracil bases prior to amplification [9–11]. However, for 5-methylcytosine (5mC) in CpG dinucleotides, deamination by formalin would be converted directly to thymine instead of uracil [3,8]. This second class of formalin artefact is not corrected by the repair treatment therefore, downstream bioinformatics approaches are necessary to attempt their removal [5].

Mutational signatures derived from whole genome sequencing (WGS) data characterise the mutational processes that have acted upon the DNA within a sample [12,13], and they hold tremendous potential for diagnosis and therapeutic guidance [14–18]. Single base substitution (SBS) signatures are derived by considering the type of specific base pair change (e.g. C>T or C>A, *etc.*) together with the flanking base pair context (e.g. ACA>ATA, or ACA>AAA, *etc.*) [12,13]. The recently updated mutational signature catalogue provides a comprehensive source of mutational processes active in human cancers that is derived from an unprecedentedly large number of samples [19]. As the artefactual mutations from FFPE preservation will bias mutational profiles, they have to be taken into account when unravelling mutational processes from FFPE samples.

Here, we use the statistical machinery of mutational signature analysis to derive mutational footprint caused by formalin exposure during FFPE biospecimen processing. First, we identify the “formalin artefact” mutational signatures in both unrepaired and repaired FFPE samples, using paired FFPE and FF sequencing data from the same samples. We next design and validate a decomposition algorithm, FFPEsig, to subtract FFPE artefacts and thereby infer mutational profiles of biological origin in genome sequencing data from an FFPE specimen. Our method enables robust mutational profile correction of FFPE samples for research and potential clinical implementation.

## Results

### Mutational signatures of formalin fixation

#### Formalin fixation artefacts are predominantly C>T mutations

To identify artefacts signatures, we used publicly available targeted panel sequencing data from two previous studies [8,11], in which triplicate samples (FFPE-repaired, FFPE-unrepaired and FF) were available. The study by Prentice *et al.* (hereafter study 1) comprised colorectal cancers (*n*=3), and each cancer included nine samples: one FF sample, four unrepaired and four repaired FFPE samples that were sequenced after a fixation time of 2, 15, 24 and 48 hours respectively. In addition, study 1 included patients (*n*=29) for whom repaired and unrepaired FFPEs were available. In the study by Bhagwate *et al.* (hereafter study 2), triplicate samples from benign breast tissue (*n*=4) were available. In total, we obtained 110 FFPE samples, of which 32 (29%) had matched FF (see Methods & Materials).

We first focused on samples with matched FF available, and examined the set of mutations detected in FFPE samples but not detected in matched FF samples (termed FFPE-only or discordant mutations). Within the study 1 sample set, we discovered that C>T discordant mutations were common (45.8% and 21.1% in unrepaired and repaired samples, respectively). T>C mutations were also common (53.5% and 76.3% in unrepaired and repaired FFPEs, respectively; Supplemental Fig 1). Discordant FFPE-only mutations from study 2 also tended to be C>T mutations (98.9% in unrepaired and 76.6% in repaired FFPEs), but very few T>C mutations were detected in this second study (0.55% in unrepaired and 11.6% in repaired FFPEs; Supplemental Fig 2).

To examine whether T>C mutations were true artefacts of FFPE, we counted the proportion of C>T and T>C mutations present in two or more of the set of samples from a patient (‘concordant mutations’). On average, about 30% C>T mutations were shared by at least two samples, in contrast to 88% for T>C mutations (Supplemental Fig 3a). We next compared frequencies of concordant mutations between all sample-pairs across three patients: 12% of C>T mutations and 59% of T>C mutations were shared by one sample-pair on average (Supplemental Fig 3b). Furthermore, C>T discordant mutation loads increased with formalin fixation time in both repaired (slope=0.80, intercept=89.68) and unrepaired FFPE samples (slope=7.48, intercept=164.81) (Fig 1a). However, the T>C discordant mutation loads decreased with fixation time in unrepaired FFPE (slope=-0.63, intercept=350.85), but increased in repaired FFPEs (slope=1.02, intercept=364.62) (Supplemental Fig1). Taken together, our results suggested that C>T mutations are the predominant true formalin induced artefacts, and that T>C mutations are likely caused by other sources of mutational noise rather than formalin fixation.

**Fig1.**
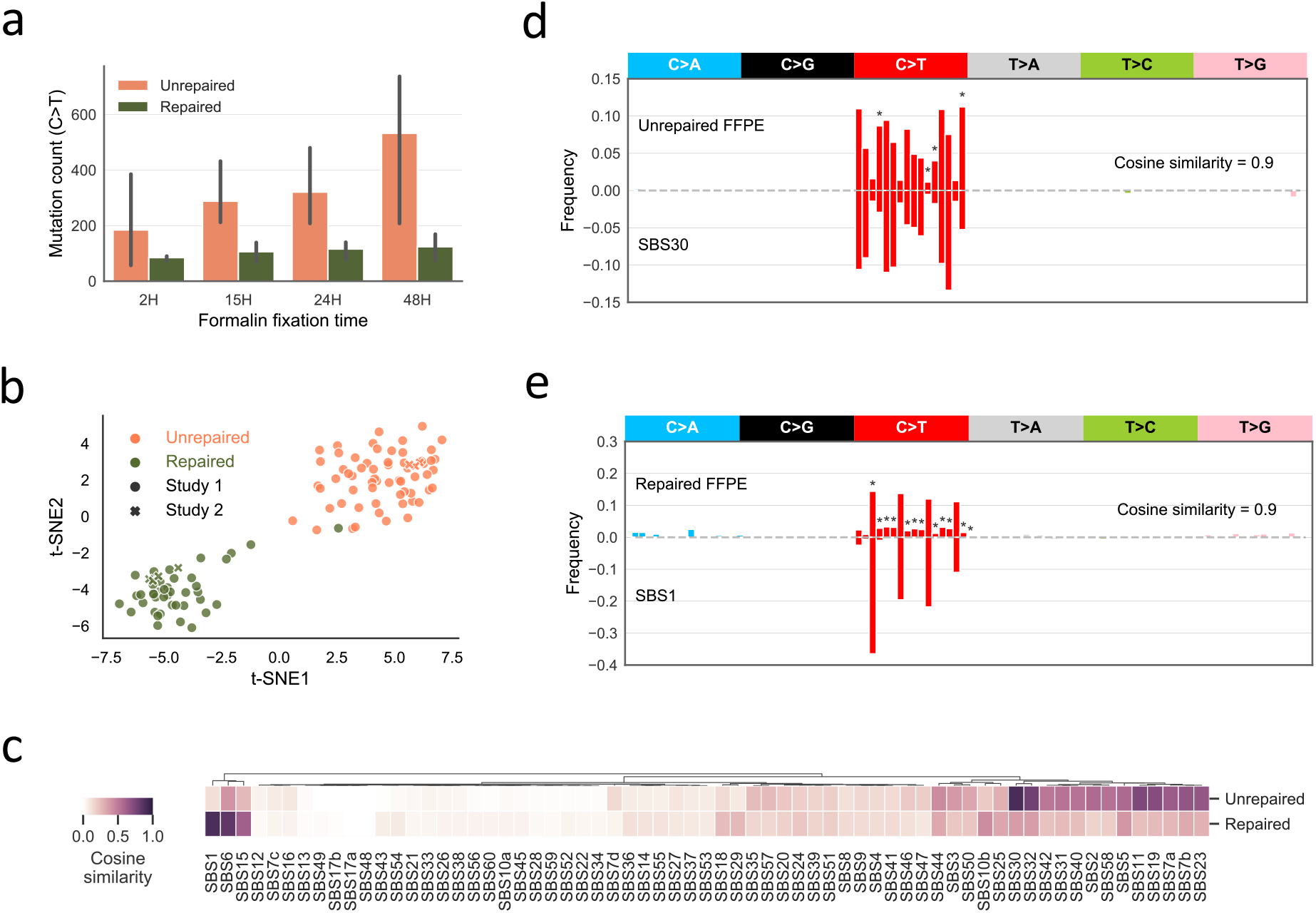
FFPE artefact signatures. (a) C>T mutation count in FFPE samples increases with formalin fixation time. We used FFPE-only C>T mutations, referring to C>T mutations that are only discovered in FFPE samples not in matched FF. The error bar shows standard deviation for measurements made on three individuals. (b) Cluster of n=110 normalised FFPE mutational profiles from two different studies [11,8]. The cluster is represented by t-SNE on cosine metric of normalised 80-channel (without T>C) FFPE-only mutational profiles (see Methods & Materials). Each FFPE sample is classified as unrepaired (with UDG treatment; pink dots) or repaired (without UDG; green dots). The two studies are marked using circle or cross shape. (c) Comparison of FFPE signatures to COSMIC V3 SBS signatures. (d) Unrepaired FFPE signature is highly similar to SBS30. C>T mutation channels with fold change over 2 are marked with asterisk. (e) Repaired FFPE signature is highly similar to SBS1.

#### Unrepaired formalin signature is highly similar to SBS30; repaired formalin signature is highly similar to SBS1

We next used all FFPE-only mutations (T>C excluded) to learn FFPE signatures. Analysis was performed on all FFPE samples (n=110). The samples in the respective studies were sequenced using different cancer gene panels, thus the ‘mutational opportunities’, determined by the frequency of each trinucleotide context in the panel, differed between studies (Supplemental Fig 4). Therefore, we applied the study-specific normalisation on the mutation counts to enable direct comparison between the studies (see Methods & Materials). The cluster of normalised mutational profiles from the entire combined set of n=110 FFPE samples was represented using t-distributed stochastic neighbour embedding (t-SNE) [20] (Fig 1b). Samples from the two studies showed no batch effect and clearly separated into two clusters of unrepaired and repaired samples. A single repaired sample from study 1 clustered with unrepaired FFPEs, which we suspect is due to poor response to UDG treatment [21]. In addition, we clustered T>C mutational profiles after normalisation, but discovered a clear batch effect and found no consistent error patterns (Supplemental Fig 3c).

To exclude possible outliers, we used t-SNE clustering to select representative samples. We performed an iterative process where each iteration was defined by the random seed inputted to the t-SNE algorithm. For each t-SNE embedding, we calculated the spatial density of the clustered data measured by a gaussian kernel, and selected samples in regions of high density (density>0.018) as our representative sample subset (Supplemental Fig 5a). The averaged values of all mutation channels from this representative subset generated one set of FFPE signature candidates. Our final FFPE signatures were derived from the mean of 100 candidates collected from 100 t-SNE embeddings (Supplemental Fig 5b and 5c; Supplemental Table 1).

We then compared the derived FFPE artefact profiles to the latest COSMIC SBS signatures (V3 - May 2019) [19] (Fig 1c), and found that unrepaired and repaired FFPE signatures are highly similar to SBS30 and SBS1 respectively (cosine similarity 0.90 for both; Fig 1d and 1e). SBS30 has been validated as a mutational footprint of *NTHL1* mutations that disrupt base excision repair (BER) [22,23]. SBS1 is well-known as a ‘clock-like’ signature that positively correlates with patient age, as a consequence of spontaneous deamination of methylcytosine [24]. We note that the unrepaired FFPE signature shared even greater similarity with COSMIC V2 (March 2015) signature 1 (0.95), which was inferred from a smaller cohort compared to SBS1 of V3.

Despite the high similarity, there were certain mutation channels that differed between FFPE signatures and the two known mutational processes. We marked the mutation channels if the fold-change was over 2 (Fig 1d and 1e). Unrepaired FFPE signature differs in NCT context. The repaired FFPE signature mostly differs in non-CpG mutation channels which are absent in SBS1(V3) but present in sig 1 (V2). Those small proportions of mutations in non-CpG channels of repaired FFPE signature are likely due to the artefactual mutations escaped from the UDG repairing process.

### Development and validation of FFPE artefacts correction algorithm using synthesised data

We designed and implemented an algorithm we called “FFPEsig” to correct artefacts from FFPE mutational profiles (see Methods & Materials). The algorithm decomposes the observed aggregate mutational catalogue of one given FFPE sample as the combination of FFPE-artefacts and the true biological mutations. To test the performance of the method, we added FFPE-artefacts to all PCAWG samples *in silico*, and then attempted to remove these artefacts using FFPEsig [19,25]. Fig 2a shows the true, simulated and corrected profiles for one colorectal cancer (CRC) sample. In this case, FFPEsig successfully inferred the biological mutation catalogue with ~0.99 accuracy, measured by cosine similarity on C>T channels. The correction accuracy was slightly higher when we used the full 96-channel (Supplemental Fig 6), but the predominance of formalin associated mutations in the C>T channels meant the gain was minimal. Therefore, hereafter we evaluated our correction accuracy focusing only on C>T mutation channels.

**Fig2.**
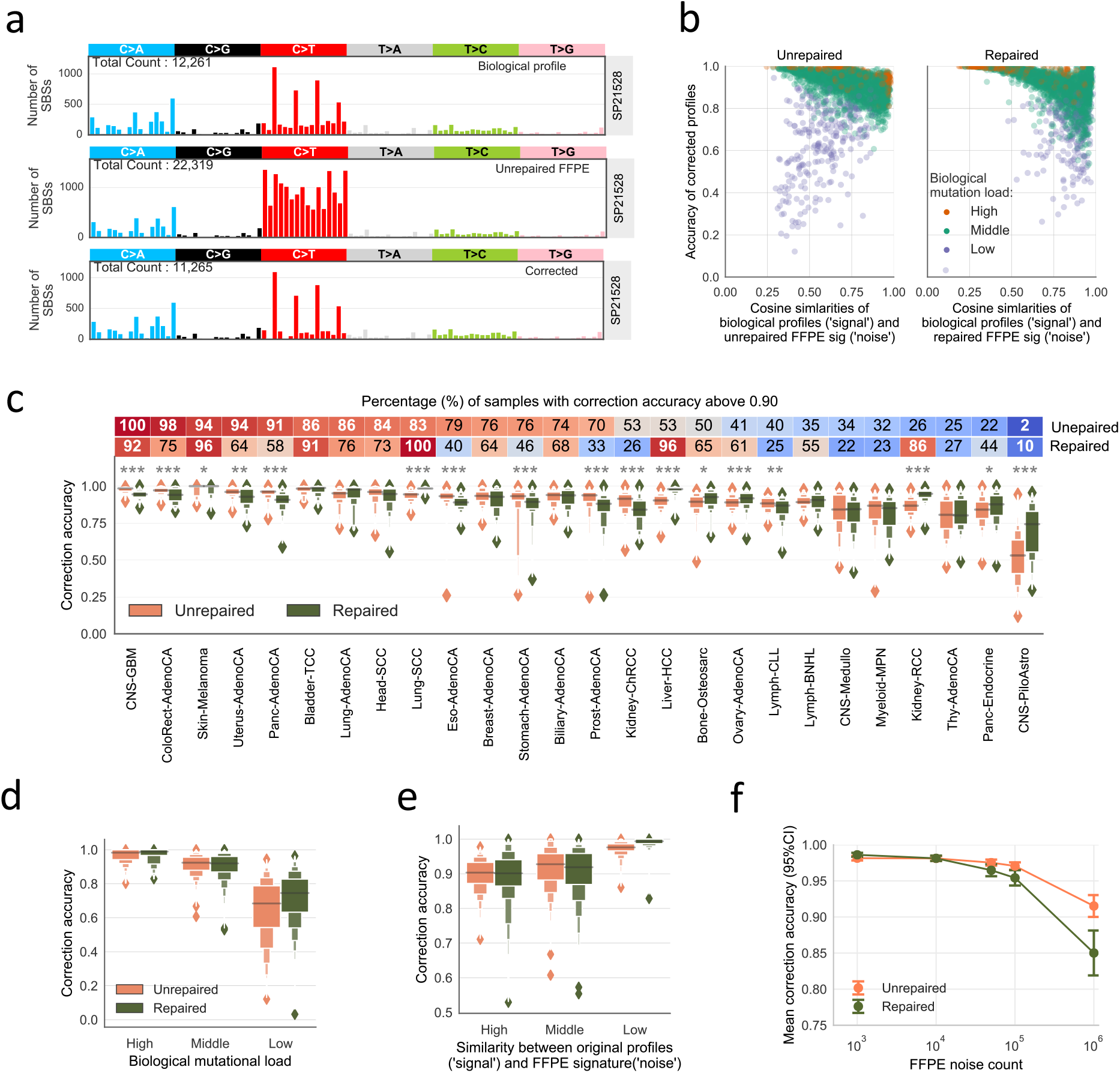
Correction of FFPE artefacts in synthetic FFPE samples. We added FFPE noise signatures to biological mutational profiles in PCAWG dataset [20] to simulate FFPE mutational profiles. (a) Artefacts correction result of one colorectal cancer (SP21528). From top to bottom, the three panels are original/biological mutational profile, simulated FFPE mutational profile (unrepaired) and the corrected somatic mutation catalogue. (b) Correction accuracy for all simulated data. The left panel shows results for unpaired FFPEs and the right panel is for repaired FFPEs. The x-axis shows cosine similarities between original profiles (‘signal’) and the FFPE signatures (‘noise’). We also group the data into three categories according to the biological mutation load, namely high (top 10%, orange dots), low (bottom 10%, purple dots) and middle (the remaining ones, green dots). (c) Correction accuracy for different cancer types. Cancer types with at least 20 samples are used here. The difference between unrepaired and repaired FFPE correction accuracy is shown above each box-pair using two-sided Mann-Whitney *U* test. *P* value <= 0.001 (***); 0.001 < *p* value <= 0.01 (**); 0.01 < *p* value < =0.05 (*); *p* value > 0.05 (none). The percentages of well-corrected samples (accuracy > 0.9) are annotated in the top colour bars. (d) Correction accuracy positively correlates with biological mutation load. (e) Correction accuracy negatively correlates with with similarities between ‘signal’ and ‘noise’. The three categories are use high (top 10%), low (bottom 10%) and middle (the remaining ones). (f) Correction accuracy drops with increasing FFPE artefacts in both types of FFPEs. We selected cancer types with at least 80% well-corrected samples in both unrepaired and repaired FFPEs from (c). The results are collected from simulated samples added with five different noise levels from 103, 104, 5×104, 105 to 106. The 95% confidence interval of each mean correction accuracy is marked using error bar here.

Overall, FFPEsig achieved 0.89 mean correction accuracy for both unrepaired (95% CI: 0.885, 0.893) and repaired FFPEs (95% CI: 0.887, 0.894) (Fig 2b and 2c). To examine the possible factors which could influence the artefact correction, we evaluated 1) biological mutation count; 2) the similarities between the artefact signature (the ‘noise’) and the true biological mutation catalogue (the ‘signal’). Poorly corrected cases were due to low mutation load and/or high similarity of patterns shared between the noise and signal (Fig 2b). We noticed that samples with low biological mutation load were difficult to correct regardless of how different the mutation patterns are from the FFPE signatures (purple dots in Fig 2b). We further separated these two factors and confirmed that higher biological mutation burden led to more accurate correction (Fig 2d), as well as high dissimilarity between the signal and the noise (Fig 2e; cases with low mutation load excluded).

We continued our *in silico* evaluation by examining correction performance across cancer types for simulated unrepaired and repaired FFPEs within each cancer type (Fig 2c). The efficacy of correction varied significantly across 26 cancer types. FFPEsig was most accurate in skin melanoma (mean: 0.98) due to its high mutation load (96,361 SBSs) and low similarity to the noise signatures (0.55) for both FFPE samples, followed by bladder transitional cell carcinoma (Bladder-TCC, 0.97) and lung squamous cell carcinoma (Lung-SCC, 0.96). In contrast, FFPEsig performed poorly for pilocytic astrocytoma (CNS-PiloAstro, 0.61), thyroid adenocarcinoma (Thy-AdenoCA, 0.80) and medulloblastoma (CNS-Medullo, 0.82), because of the low averaged mutation loads (from 112 to 602) and relatively higher similarity to the noise signatures (0.69-0.74) in these cancer types.

We also noticed that the algorithm had different performance between unrepaired and repaired FFPEs within certain cancer types. There were 17 out of 26 cancer types with detectable difference in correction efficacy (*p*-value < 0.05) and 12 of 17 with a highly significant difference (*p*-value < 0.001). For instance, the correction worked much better in unrepaired FFPEs for colorectal (ColoRect-AdenoCA) and pancreatic adenocarcinoma (Panc-AdenoCA), with 98% and 92% of well-corrected samples for unrepaired FFPEs respectively, in contrast to only 71% and 51% respectively for repaired ones. Since the mutation burdens were the same for two types of FFPEs within a cancer type, the significant difference is caused by true mutations being more dissimilar to the FFPE-artefact profile in unrepaired FFPEs (cosine similarity 0.49 for CRCs and 0.59 for pancreatic cancers), whereas the repaired-FFPE mutational signature was very similar to the true mutational profile (cosine similarity 0.89 and 0.90 colorectal and pancreatic cancer respectively). By contrast, FFPEsig worked successfully in repaired-FFPEs for Lung-SCC and liver hepatocellular carcinoma (Liver-HCC), with 100% and 96% well-corrected samples for the opposite reason.

Finally, we explored how the accuracy of FFPE-artefact removal changes with increasing noise of FFPE artefacts (Fig 2f). We selected four cancer types with 80% or more well-corrected samples in both repaired and unrepaired FFPEs, including 219 tumour samples from CNS-GBM, Skin-Melanoma, Bladder-TCC and Lung-SCC (Fig 2c). As expected, as the burden of artefactual mutations was increased, the correction accuracy dropped from 0.97 to 0.86 in unrepaired FFPEs, and from 0.98 to 0.84 in repaired FFPEs. Overall, FFPEsig performed equally well in both types of FFPE with up to 10^5^ noise (mean accuracy > 0.94), but its performance dropped dramatically for samples with 10^6^ noise (0.84-0.86). Thus, our method works for samples that hold reasonable signal-to-noise ratio, but not for the extreme cases, e.g. samples with 10^6^ noise in this experiment with signal-to-noise ratio around 0.0088.

### A case study of correcting FFPE artefacts in WGS FFPE CRC blocks shows consistent results with simulated data

Next, we performed whole genome sequencing on two tumour FFPE samples (unrepaired versus repaired), and on the normal tissue DNA as matched normal from the same CRC patient (see Methods & Materials; FF material was not available). The mean coverages of the sequencing data were 46X (unrepaired FFPE), 43X (repaired FFPE) and 43X (normal sample), with 98.81% or more of reads mapped to the genome (Supplemental Table 2). Following filtering (see Methods & Materials), we detected 13,208 and 6,107 somatic single base substitutions in unrepaired and repaired FFPE, respectively (Supplemental Fig 7a and 7b). In particular, the two types of dominant mutations in our FFPE samples were C>T and T>C, and together they contributed 64.7%-66.6% to the total mutations (Supplemental Fig 7b). For C>T mutations, we expected them to be a mixture of FFPE artefacts and real biological mutations, because of the relative preponderance (~35%) of C>T mutations in PCAWG CRCs. T>C mutations accounted for 41.2% and 39.8% in our unrepaired and repaired FFPEs, but only ~16% in PCAWG CRCs (Supplemental Fig 7c). Similarly, large proportions of T>C mutations were also detected in FFPE samples in study 1 (Supplemental Fig 1). As noted above, these presumably artefactual T>C mutations did not show consistent patterns (Supplemental Fig 3c). Therefore, we excluded T>C mutations from further study.

Since matched FF was not available to provide the ground truth mutational signature, we were inspired by results found in study 2 [8], where both repaired and unrepaired FFPE samples contained the majority of the variants found in the matched FF sample. Thus, we used concordant mutations with more strict filtering (variant supporting reads ≥ 5 in both FFPEs) as an approximation for the true biological mutation profile of the tumour: this yielded a total of 1040 filtered concordant mutations (Supplemental Fig 7a and 7b), and 656 of them remained after excluding T>C mutations (top panel of Fig 3a).

**Fig3.**
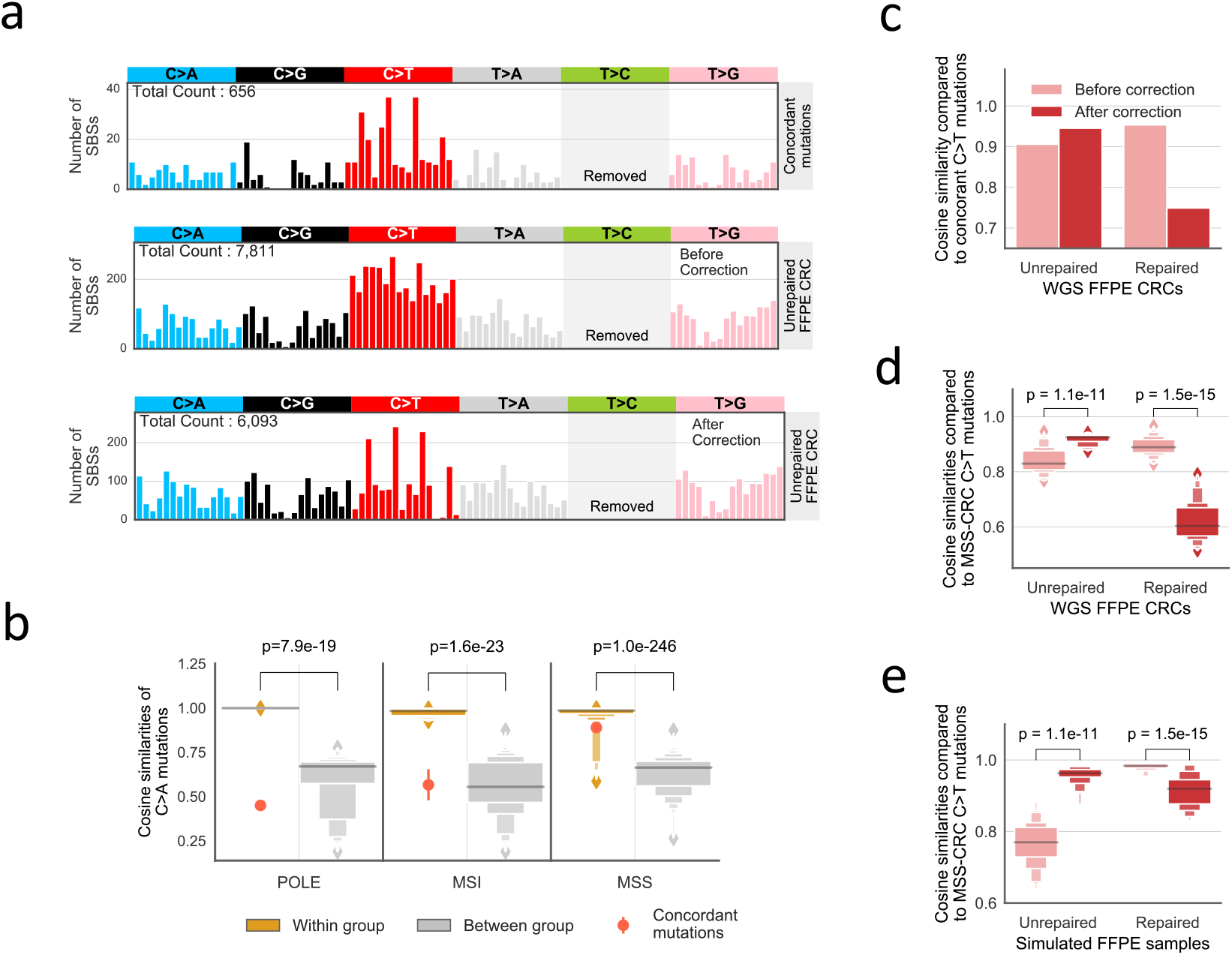
A case study of applying FFPE artefact correction method on two WGS CRC FFPE samples. The two FFPE samples are from the same CRC patient. One of FFPEs is unrepaired and the other one is repaired. (a) Correction result for the unrepaired FFPE sample. The three panels are concordant mutation catalogues (top), unrepaired FFPE CRC profiles before correction (middle) and after correction (bottom). Concordant mutations refer to variants are shared between repaired and unrepaired FFPEs with at least 5 reads supporting the variant, and their profile is taken as an approximation of true mutational catalogue of the tumour. We removed T>C mutations to show clear pattern of other mutation channels due to their large numbers. (b) Concordant C>A mutation profile is highly similar to MSS-CRC C>A mutation patterns. PCAWG CRCs are grouped based on their known labels, namely POLE, MSI and MSS. The sample-pair cosine similarities of C>A mutation patterns within and between subgroups are shown in orange and grey box plot, respectively. The *p*-values of differences for each subgroup are shown above each box-pair using two-sided Mann-Whitney *U* test. The error bar shows standard deviation.(c) Comparing correction results of two FFPE samples to concordant mutations. As the correction acts on C>T mutation channels, we compared the cosine similarity changes of original profile (pink colour) and corrected profile (red colour) on C>T channels. (d) Comparing correction results of two FFPE samples to MSS-CRCs. (e) Comparing correction results of simulated MSS-CRC FFPE profiles. We compared each simulated MSS-CRC FFPE sample to all other MSS-CRC profiles but their real biological profile to treat them the same way our WGS FFPE samples, for which the FF sample is not available.

To obtain more general knowledge about the biological mutation profiles of CRCs, we performed hierarchical clustering on the 60 PCAWG CRC samples and discovered the samples share highly homologous mutational profiles within each subtype, namely MSS, MSI and POLE (Supplemental Fig 8a). The averaged sample-pair cosine similarity is 0.90 for MSS-CRCs, 0.92 for MSI-CRCs and 0.96 for POLE-CRC, but profiles between subtypes are significantly different (Supplemental Fig 9a). To identify the most “conserved” mutation patterns within each subtype, we performed a similar analysis on six mutation types separately, which showed C>A and C>T mutations have the strongest power in classifying CRC subtypes (Supplemental Fig 8b and 9b). Therefore, we compared the concordant C>A mutations observed in our case to the PCAWG CRCs and identified that our sample was a MSS-CRC (Fig 3b).

We next applied FFPEsig on the observed mutation counts from the two FFPE samples and valuated the corrected profiles (Fig 3a and Supplemental Fig 10) by comparing them to concordant mutation catalogue as well as all PCAWG MSS-CRC samples, under the assumption that after removing artefacts the mutational profile of our samples should show higher similarity to both ‘positive controls’. For unrepaired FFPE CRC, the accuracy improved from 0.906 before correction to 0.945 after correction to concordant mutations (Fig 3c). When compared to MSS-CRCs, the correction led to a significant increase in cosine similarity from 0.841 to 0.918 (Fig 3d). However, correction on repaired FFPE CRC generated the opposite results (Fig 3c and 3d). We validated our observations using simulated FFPE MSS-CRCs and confirmed that the correction was only beneficial for unrepaired not repaired FFPEs (Fig 3e). This was because the biological MSS-CRC profiles are highly similar to the repaired FFPE signature (0.98 on C>T channels) and so our correction method could not distinguish true mutations from artefacts.

We further investigated how our corrected profile from unrepaired FFPE could contribute to CRC subtyping. Application of MSIsensor [26] detected 8.3% of microsatellite sites with somatic changes in the unrepaired FFPE sample, but only 0.23% from the repaired FFPE. 8.3% exceeds the 3.5% threshold to call MSI [26], and so application of MSIsensor to an unrepaired FFPE sample could lead to miscalling of MSI status. We therefore attempted to classify the sample using the ‘conserved’ mutation patterns within CRC subtypes (described above). The unrepaired FFPE sample was equally similar to both using observed C>A and C>T trinucleotide mutational counts together or only C>T mutations (Supplemental Fig 11a and 11b). However, following correction using FFPEsig, we could clearly distinguish that the sample was MSS. In addition, we found that the C>A mutation pattern itself could also classify our sample (Supplemental Fig 11c). As FFPEsig mostly in C>T channels, C>A patterns were almost the same with or without correction (0.99).

### Potential of using 80-channel signatures for refitting analysis in FFPE samples

T>C were common in some but not all FFPE samples in our dataset, and perhaps resulted in differences in sequencing library preparation methodology between studies. To attempt to control for this unexplained variation, here we examined the impact of removing all T>C variants during signature refitting analysis. We compared the attributed mutation count (or activity) of each signature by supplying our refitting model with 80-channel (80c; T>C removed) and 96-channel (96c) signatures on PCAWG mutational catalogues (see Methods & Materials; Supplemental Fig 12). The log_10_ signature activity ratio of 80c to 96c was used to estimate how consistent both results were, and we termed this value as an inconsistency rate. The bigger the absolute inconsistency rate is, the more different the attributions are.

We refitted 10,312 mutational signature activities for 29 active signatures from 2,726 PCAWG genomes (Fig 4a), and an additional 54 genomes were excluded from original PCAWG dataset due to either low reconstruction accuracy (<0.85; n=35) by 96c signatures or too small of a sample size (<10 cases per signature per cancer type; n=19). The mean inconsistency rate among 10312 refits was 0.013 (95% CI: 0.0076, 0.1783) (middle panel of Fig 4a). We considered signatures with inconsistency rate between −0.30 to 0.18, equivalent to actual activity ratio from 0.5 to 1.5, as having well-refitted results. Of the originally inferred 10312 signature activities that used 96c data, 8938 (86.7%) were well-refitted when only 80c data was used. 24 of 29 signatures were considered well-refitted.

**Fig4.**
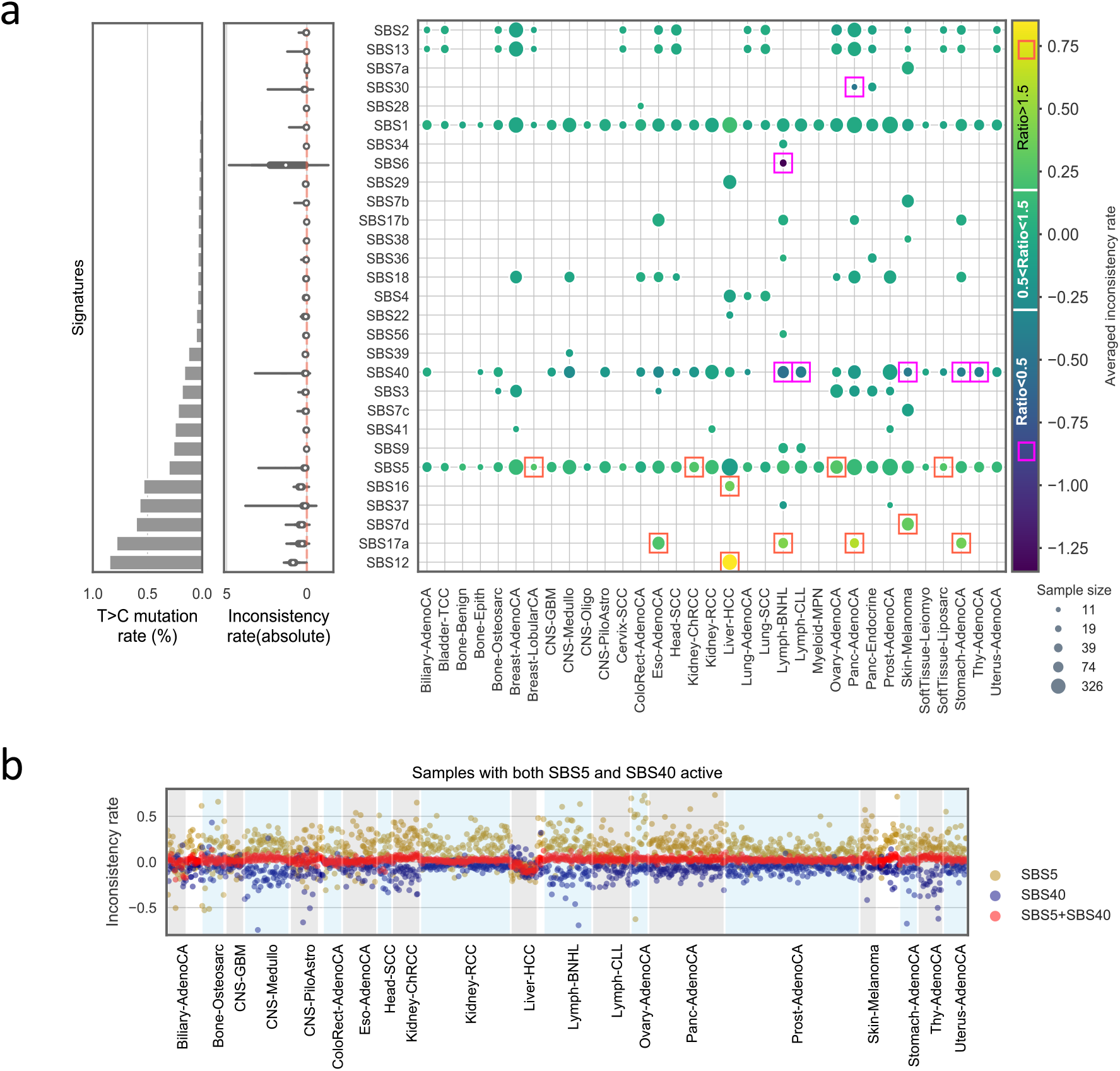
Comparison of signature activities inferred by signatures with and without T>C mutations. We inferred signature activities using 96-channel (96c) and 80-channel (80c; without T>C) signatures on PCAWG mutational profiles. Here we use inconsistency rate as a measurement for how well the inferred activities agree with each other. Inconsistency rate is calculated as log10(activity_80c/activity_96c). (a) Activities inferred by 96c and 80c signatures are consistent for majority of signatures. Left panel: sum of mutational probabilities in T>C channels for each signature. Middle panel: violin plot of absolute inconsistency rate for all signatures. Right panel: heatmap of mean inconsistency rate for all signatures in different cancer types. Orange rectangle marks the average activity ratio (activity_80c/activity_96c) above 1.5 (~0.18 on log10 scale), which means 80c activity is bigger than 1.5 times of 96c activity. The purple rectangle marks the averaged activity ratio below 0.5 (~ −0.30 on log10 scale), which means 80c activity is smaller than 50% of 96c activity. The radius of each circle represents the sample size (in log scale). (b) Activity flows between two similar signatures (SBS5 and SBS40). The inconsistency rates for SBS5 in all samples are in golden dots, and those for SBS40 are in blue dots. The inconsistency rate for the sum activity of SBS5 and SBS40 is shown in red dots.

For the five signatures that were poorly refitted using 80c, four of them had high T>C mutation rates, namely SBS7d, 12, 16 and 17a (left panel of Fig 4a). The inconsistency rate was significantly correlated with T>C mutation rate of signatures (Spearman’s rho=0.54, p<e-10). We grouped the refitted data based on cancer types (right panel of Fig 4a) and discovered the majority of the above five signatures with inconsistent refits were each only reported in one cancer type, except for SBS17a which was present in four cancer types. SBS6 also had a high inconsistency rate and was mostly detected in non-Hodgkin lymphoma (lymph-BNHL), likely due to the higher similarity shared with SBS1 (0.77). Taken together, removing T>C mutations had a very minor impact on refitting analysis for the majority of the cases (86.7%), apart from the minority of cases with a high T>C mutation rate.

In addition, SBS5 and SBS40 showed noticeable differences between 96c and 80c fits in several cancer types. With the knowledge of these two ‘flat’ signatures are highly similar (0.83 using 96c; 0.86 using 80c), the model could have problems distinguishing them using either 80c or 96c. Thus, we suspected that the inferred signature activity of SBS5 or SBS40 could vary individually within a sample, but the sum of the activity of the two signatures would be fairly constant. We tested our hypothesis on samples with both signatures active (Fig4b). As expected, the sum of activities converged well with the mean inconsistency rate of 0.02 (95% CI: 0.019, 0.023), but individual attribution for SBS5 was higher by 80c (mean inconsistent rate of 0.15; 95%CI: 0.14, 0.16) and lower for SBS40 (mean inconsistency rate of −0.19; 95%CI: −0.21, −0.16), and the two individual attributions were negatively correlated (Spearman’s rho=-0.69, p=6.22e-164).

Finally, we examined signatures where removal of the T>C mutations was most likely to be detrimental for signature identification. We compared all possible signature pairs among 65 COSMIC V3 SBS signatures (Supplemental Fig 13). As expected, the overall similarities between any two signatures tended to increase, especially for the originally dissimilar (<0.2) signatures pairs (Supplemental Fig 13a and 13b). Five signature-pairs became highly similar (>0.8) using 80c. Three out of them are reported to be biological/non-artificial mutation processes, namely SBS3-SBS5, SBS40-SBS12 and SBS40-SBS16 (Supplemental Fig 13c). However, two signature-pairs became even more distinguishable using 80c (Supplemental Fig 13c). Therefore, we concluded that reducing to 80 channel signatures by removal of T>C channels tended to have a minor effect on signature identification.

## Discussion

In this study, we derived genome-wide mutational signatures that result from formalin exposure in FFPE biospecimens and designed an algorithm, FFPEsig, to detect and remove artefactual-FFPE mutations from measured mutational profiles. The accuracy of FFPEsig was demonstrated on synthetic FFPE samples. Accuracy was generally very high. We note poorer performance occurred when (a) biological mutation loads were low and (b) for samples where the true mutational profile closely resembled the FFPE-artefact signature - we note these circumstances are straightforward to identify in practice and so it is clear when FFPEsig can be safely applied. We note that the statistical machinery within FFPEsig is generalisable, and could be repurposed to correct for “mutational noise” from any source.

The repaired FFPE signature discovered in this study is highly similar to the aging signature SBS1 (Fig 1e). Both formalin-mutagenesis and the process leading to biological SBS1 are caused by deamination of 5-methylcytosine (5mC) (SBS1 is due to spontaneous deamination *in vivo* whereas the FFPE signature is caused by chemical deamination *in vitro* [5,24]). Unfortunately, this high similarity precludes the study of the activity of the aging signature in repaired FFPEs, which is active in all tumour genomes [24]. Similarly, the signature associated with unrepaired FFPE samples is highly similar to SBS30 and therefore would also distort the study of SBS30 in FFPE samples (Fig 1d). However, biological SBS30 occurs more rarely: it is caused by loss-of-function in glycosylases in BER due to biallelic inactivation mutations in *NTHL1*, and patients carrying this variant are diagnosed as *NTHL1* tumour syndrome with an increased lifetime risk for CRC, breast cancer, and colorectal polyposis [22,23,27]. More generally, our results show that there is not necessarily a direct 1-to-1 mapping relationship from mutational process to a unique signature profile (as also questioned in [28]) as distinct mutational sources can cause similar profiles. Nevertheless, our findings speak to the utility of constructing a common carcinogen signature database [28,29].

The accumulation speed of C>T artefacts in unrepaired FFPEs suggests that UDG “repair treatment” rectified DNA deamination damages to a large extent (Fig 1a). Therefore, fixation time is an important pre-analytical factor of determining the burden of FFPE-artefact mutations, which could influence the downstream signature analysis. Further, large numbers of putatively artefactual T>C mutations can be present in FFPE samples and biological interpretation of these must be performed with extreme care. Indeed, Marchetti *et al.* identified 22 out of 24 (92%) previously reported ‘novel’ mutations in *EGFR* to be FFPE artefacts, and those 22 mutations were either C>T or T>C [30]. So far, we have not found evidence showing which chemical agent in formalin could cause deamination of adenine, as this would result in hypoxanthine residues and further preferentially pair with cytosine to generate A:T>G:C artefacts [31]. However, regardless of the unclear mutagenic mechanism, once the wrong residuals were generated on the DNA, multiple PCR amplifications of very small amounts of DNA from paraffin-embedded tissues would make the artefacts easily observed from the data [30].

## Conclusion

In conclusion, here we identified two mutational signatures, linked to repaired and unrepaired FFPE, which are highly similar to COSMIC signatures SBS1 and SBS30, respectively. We further developed FFPEsig software to accurately remove FFPE-induced mutational artefacts and demonstrated efficacy *in silico* and in new samples. Careful application of our approach will enable the robust study of mutational signatures in the enormous FFPE archives that exist around the world.

## Methods & Materials

### Targeted sequencing data

We used targeted sequencing data from two previous publications [8,11]. Prentice *et al.* has collected three groups of samples from CRC patients, namely fixation, baseline and blockage, to examine the impact of three factors on somatic mutation detection in clinical FFPE samples [11]. The three factors were formalin fixation time (fixation; n=3), DNA extraction kits (baseline; n=20) and storage time (blockage; n=9). Samples collected in the fixation group were fixed in formalin for 2, 15, 24 and 48 hours for both repaired and unrepaired FFPEs, and paired FF samples were also available. To validate if true somatic mutations are detectable in FFPE samples, Prentice *et al.* applied several filters on the mutation calling results, which could have filtered FFPE artefacts out. Thus, for our purpose of learning FFPE noise signatures, we have included all data but those passed the somatic filters.

To study possible batch effects, we also included targeted panel sequencing data from study 2 in our analysis [8]. There were four normal breast tissues collected in the study. For each of them, triplicate samples were collected, fresh frozen, repaired and unrepaired FFPE. We summarised the general sample information here and more details can be found in original studies.

### Mutational opportunities for targeted sequencing data

The FASTA sequences for targeted regions for study1 were downloaded from https://www.ncbi.nlm.nih.gov/sites/batchentrez and for study2 were from https://m.ensembl.org/info/website/tutorials/grch37.html. To obtain mutational opportunities, we calculated 96-channel mutation context frequency from the second to the last second nucleotide within each sequence. We assumed one genomic location was the mutated loci and added 1 count to all mutable channels with the sequence contexts of this loci. We applied this calculation over all sequences and normalised the 96-trinucleotide counts to sum up to 1 as the mutational opportunity vector for the given targeted regions (Supplemental Fig 4a and 4b). The whole genome mutation opportunity was taken from [32] (Supplemental Fig 4c).

### Discovery of FFPE signatures

To derive FFPE signatures, we pre-processed the whole mutations list to exclude non-FFPE artefacts as much as possible. In both studies, mutations were excluded if they met any of the following criteria, 1) being detected in a matched FF sample; 2) being detected in matched normal samples; 3) with >0.9 posterior probability of being somatic mutations. The remaining mutations were used to generate 96-channel mutation counts by SigProfilerMatrixGenerator [33]. We normalised mutation counts from the two studies separately using their corresponding mutational opportunities. Specifically, the original mutation counts were divided by the mutational opportunity of the targeted regions and multiplied by mutational opportunity of whole genome context. The final normalised mutational probabilities were merged from two studies and non-T>C channels were further taken to derive FFPE signatures (Fig 1b), whereas T>C channels were analysed separately (Supplemental Fig3c).

To derive FFPE signatures, we first applied t-distributed Stochastic Neighbour Embedding (t-SNE) for dimensionality reduction for the cosine distance matrix of the merged 80-channel mutational probabilities. Based on the two principal components provided by t-SNE, we defined well representative samples for two repaired and unrepaired FFPE clusters using data point density estimated by gaussian kernel (from scipy.stats) (Supplemental Fig 5a). The high-density samples (>0.018) were used to generate one set of FFPE signature candidates. With repeating the above procedure for 100 times, we took the averaged values of each channel as the final FFPE signatures (Supplemental Fig 5b and 5c).

### Algorithm/FFPEsig for FFPE artefacts correction

We denote the observed mutation counts from the FFPE sample by *V*, which was considered as a linear combination of artefact signature *W*_1_ and biological mutation frequency *W*_2_ with their corresponding attributions/activities *H*_1_ and *H*_2_. Thus, we have:

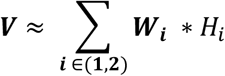

In this model, *V* and *W*_1_ were known and the task was to infer *H* = [*H*_1_, *H*_*T*_]^*T*^ and *W*_2_. Here, we utilised generalized Kullback-Leibler (KL) divergence between reconstructed 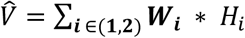 and the observed profile *V* as the cost function and applied Lee and Seung’s multiplicative update rules [34] to minimize the cost function.

The whole process of one iteration started with randomly generated initial values for *W*_2_. We then updated *H* using the multiplicative rules [34] followed by *W*, in which only *W*_2_ was updated. From the updated *W* and *H*, we got 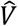. The generalised KL divergence, between *V* and 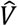 was computed and saved. This update process iterated over 200 steps by default until it met our termination criteria defined here. We calculated the convergence ratio using the average KL divergence from the last batch of 20 iterations divided by the second last batch of 20 iterations. The algorithm would terminate if the convergence ratio reaches 0.95. The maximum iteration by default was up to 3000. The above one whole process provided inferred *W*_2_ and *H* as one candidate solution. We collected 100 candidate solutions using different random seeds and averaged them as our final solution for all samples analysed for FFPE noise correction in this study.

### Simulation of FFPE samples

To simulated FFPE samples for algorithm performance validation, we added different amounts of FFPE artificial mutations with Poisson noise to biological mutation catalogues of 2,780 canner genomes provided in Pan-Cancer Analysis of Whole Genomes (PCAWG) project by International Cancer Genome Consortium (ICGC) [19,25]. The data is available to download from https://www.synapse.org/#!Synapse:syn11801889. Additionally, subtype labels of PCAWG CRC samples used in the case study were also downloaded from the same site.

### DNA extraction and genome sequence of FFPE CRCs

The male patient with ulcerative colitis was diagnosed with cancer in the transverse colon at age 48 in St. Mark’s Hospital, London, United Kingdom. Formalin-fixed paraffin-embedded (FFPE) sections of 10μm thickness were deparaffinized, rehydrated and lightly stained with methyl green. The annotated H&E was used as a guide for epithelial enrichment through targeted needle scraping of slides (for estimated epithelial cellularity >50%). To collect matched normal tissue, targeted scraping of serosal tissue from FFPE blocks was taken from a small intestinal segment distal to the cancer. DNA was extracted using a modified protocol of the High Pure FFPE DNA Isolation Kit (Roche Life Science, Penzburg, Germany). The normal tissue DNA sample and one tumour DNA sample were repaired using the NEBNext FFPE DNA Repair Mix (New England Biolabs, Inc) following the manufacturer’s recommendations. The remaining tumour DNA was left unrepaired. DNA libraries were prepared using the NEBNext Ultra II DNA Library Prep Kit for Illumina (New England BioLabs, Ipswich, Massachusetts, USA), followed by equimolar pooling strategy. Finally, all DNA libraries were sequenced on NovaSeq S2 for 50bp paired end reads.

### Somatic variants calling in WGS FFPE CRCs

The paired-end reads underwent initial quality control with FastQC [35] followed by default adaptor trimming with Skewer [36] and were subsequently aligned to GRCh38 reference genome with BWA-MEM [37]. Aligned reads were sorted by genome coordinate (SortSam, Picard) and duplicate reads were flagged with GATK’s MarkDuplicates [38]. The two FFPE tumour samples were called against the matched normal separately using the Mutect2 somatic variant caller from GATK [38]. Variants were marked with filters by FilterMutectCalls. Variants were kept if they were PASS by Mutect2, aligned to a canonical chromosome, had a total allelic depth of greater or equal to 10 in both the tumour and normal sample and had 3 or more reads supporting the alternative allele in the tumour sample. The filtered variants from two FFPE tumour samples were merged into a single VCF file using VCFtools [39].

We used Platypus on the merged VCF file as the candidate somatic variant list and integrated local alignment with multi-sample variant calling to assess the evidence for these variants across all samples [40]. The resulting VCF file was further filtered to only contain variants 1) if the FILTER flag was PASS or other acceptable filters (alleleBias, Q20, QD, SC, HapScore); 2) the variant was not a known germline variant; 3) a genotype was called for all samples; the genotype phred score was 10 or more in all samples; 4) the normal sample had no reads containing the variant and at least 3 or more reads supported the variant in a tumour sample. Variants present in two FFPE samples with 5 or more supporting reads were classified as concordant mutations.

### Signature refitting analysis

To validate if signature refitting analysis could use 80-channel spectra without T>C, we dropped T>C mutation channels of COSMIC SBS signatures and renormalised them to sum up to 1. The original activities inferred using 96-channel signatures for PCAWG cohorts were obtained from https://dcc.icgc.org/releases/PCAWG/mutational_signatures/ [19,25]. The active signatures for each sample were selected if the original activities >0. We next refitted 80c and 96c active signatures to the mutational catalogues with and without T>C mutations accordingly using our locally implemented refitting algorithm to exclude possible bias introduced by different tools. The refitting algorithm used the same multiplicative update rules and termination criteria from FFPEsig, but was different in two aspects, 1) the number of signatures was flexible which depended on the active signatures in each sample; 2) only *H* was updated in each iteration. The inferred activities for 80c-signatures were then rescaled by dividing total mutation frequencies of non-T>C mutation channels of 96c spectra. The rescaled 80c attributions were used to compare to those inferred from 96c signatures.

## Data and code access

Submission of BAM files of sequenced data to EGA is in progress. The VCF files generated in our study are available from the corresponding authors, upon reasonable request. FFPEsig is implemented in python which is available to download from https://github.com/QingliGuo/FFPEsig, as well as analysis code and data used in this study.

## Abbreviations

FFPE: Formalin fixation and paraffin embedding
FF: fresh frozen
UDG: uracil DNA glycosylase
PCAWG: Pan-Cancer Analysis of Whole Genomes
COSMIC: The Catalogue of Somatic Mutations in Cancer
SBS: single base substitutions
CRC: colorectal cancer
MSI: microsatellite instability
POLE: proofreading subunit of polymerase epsilon
MSS: microsatellite stability
EGFR: epidermal growth factor receptor
PCR: polymerase chain reaction
BAM: Binary Alignment Map file
t-SNE: t-distributed Stochastic Neighbour Embedding

## Acknowledgements

We thank Virinder Singh Reen and Dr. Ignacio Vázquez-García for helpful comments on the manuscript. We thank the team at St Mark’s Hospital London, UK, Andrea Sottoriva, Inma Spiteri, Ann-Marie Baker, Salpie Nowinski, Jacob Househam and Chris Kimberley for support with FFPE sample provision and analysis. This research utilised Queen Mary’s Apocrita HPC facility, supported by QMUL Research-IT. http://doi.org/10.5281/zenodo.438045. We also thank Prof. Amy C. Degnim, Dr. Chen Wang and the Mayo Clinic for help with data sharing.

## Author’s contributions

Q.G, T.A.G and V.M. conceived the study. Q.G. designed, carried out the data analysis, designed and implemented the algorithm and interpreted the initial results. V.M. designed the algorithm and supervised data analysis. Q.G and E.L. carried out the WGS FFPE case study. I.AB provided FFPE samples and performed genome sequencing. K.C performed mutation calling on the FFPE case. Q.G., V.M., T.A.G. and E.L. participated and contributed in results discussion and interpretation. Q.G. and T.A.G wrote the manuscript. E.L. and V.M edited the manuscript. V.M. and T.A.G supervised the project. All authors read and approved the final manuscript.

## Ethics approval and consent to participate

The archival colorectal cancer studied was collected and analysed in accordance with ethical approval from the UK Research Ethics Committee (REC: 18/LO/2051 IRAS:249008 - Fulham committee). The sample was anonymised to the researchers.

## Supplemental Figures

**Supplemental Fig1.**
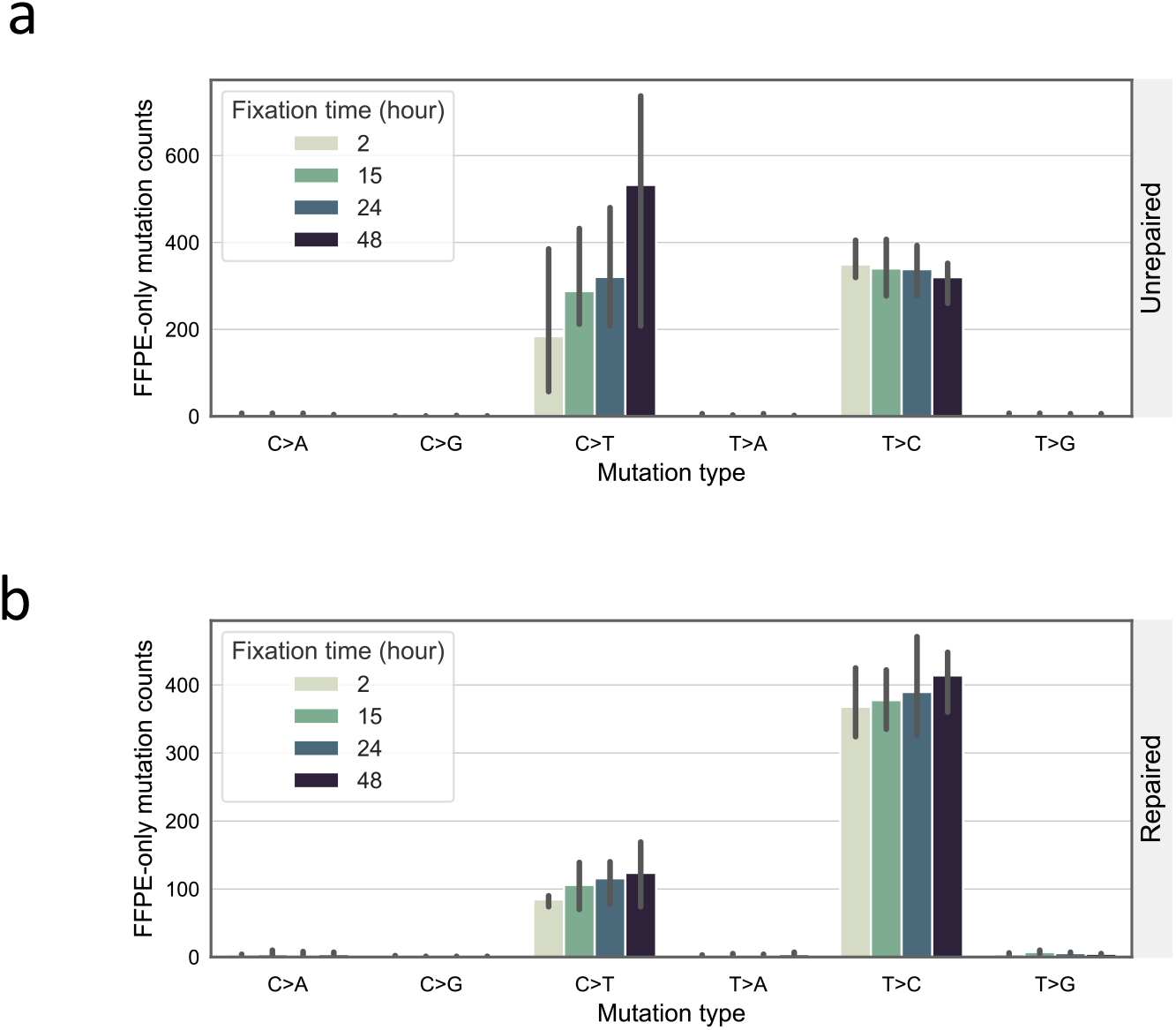
FFPE-only mutations with increasing formalin fixation time. FFPE-only mutations here refer to those are not present in matched FF sample and the data is from fixation group in study 1 [11] (see Methods & Materials). (a) Mutation count for six mutation types in unrepaired FFPE samples (without UDG treatment). For each mutation type, we show the mutation counts detected in four FFPE samples being fixed in formalin for 2, 15, 24 and 48 hours respectively. All data is collected from three patients. The error bar shows standard deviation for measurements made on three individuals. (b) Mutation count in repaired FFPE samples (with UDG treatment).

**Supplemental Fig2.**
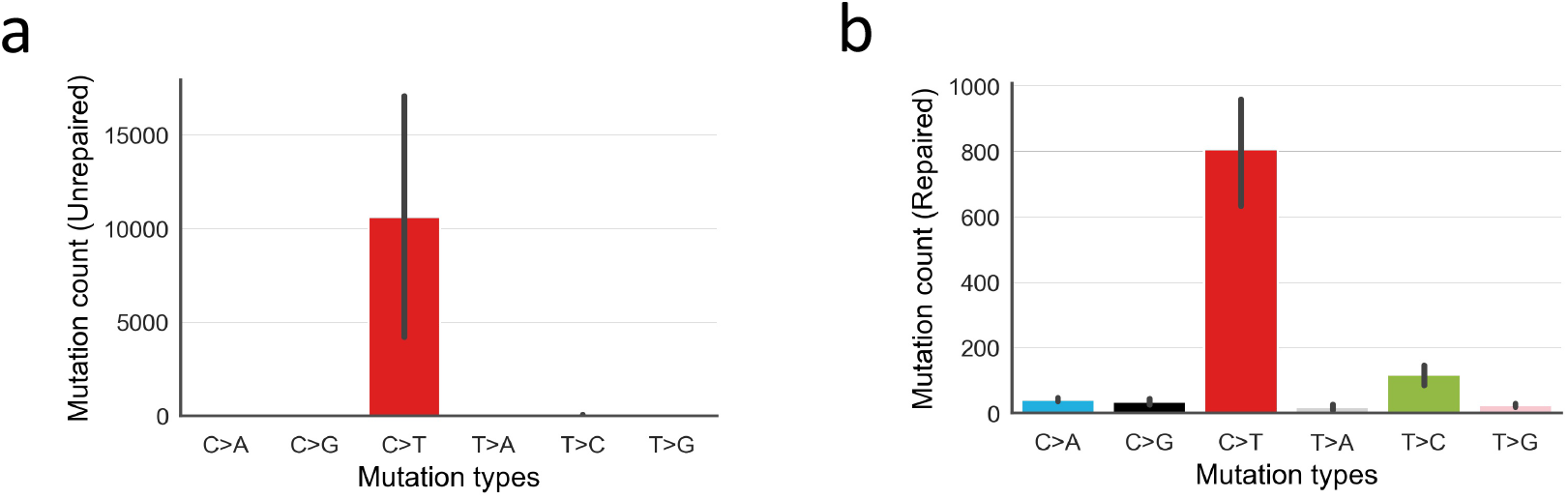
FFPE-only mutations in six basic mutation types in study 2. FFPE-only mutations here refer to those are not present in matched FF sample. The data is collected from four patients in study 2 [8] (see Methods & Materials). (a) for unrepaired FFPEs. (b) for repaired FFPEs. The error bar shows standard deviation for measurements made on four individuals.

**Supplemental Fig3.**
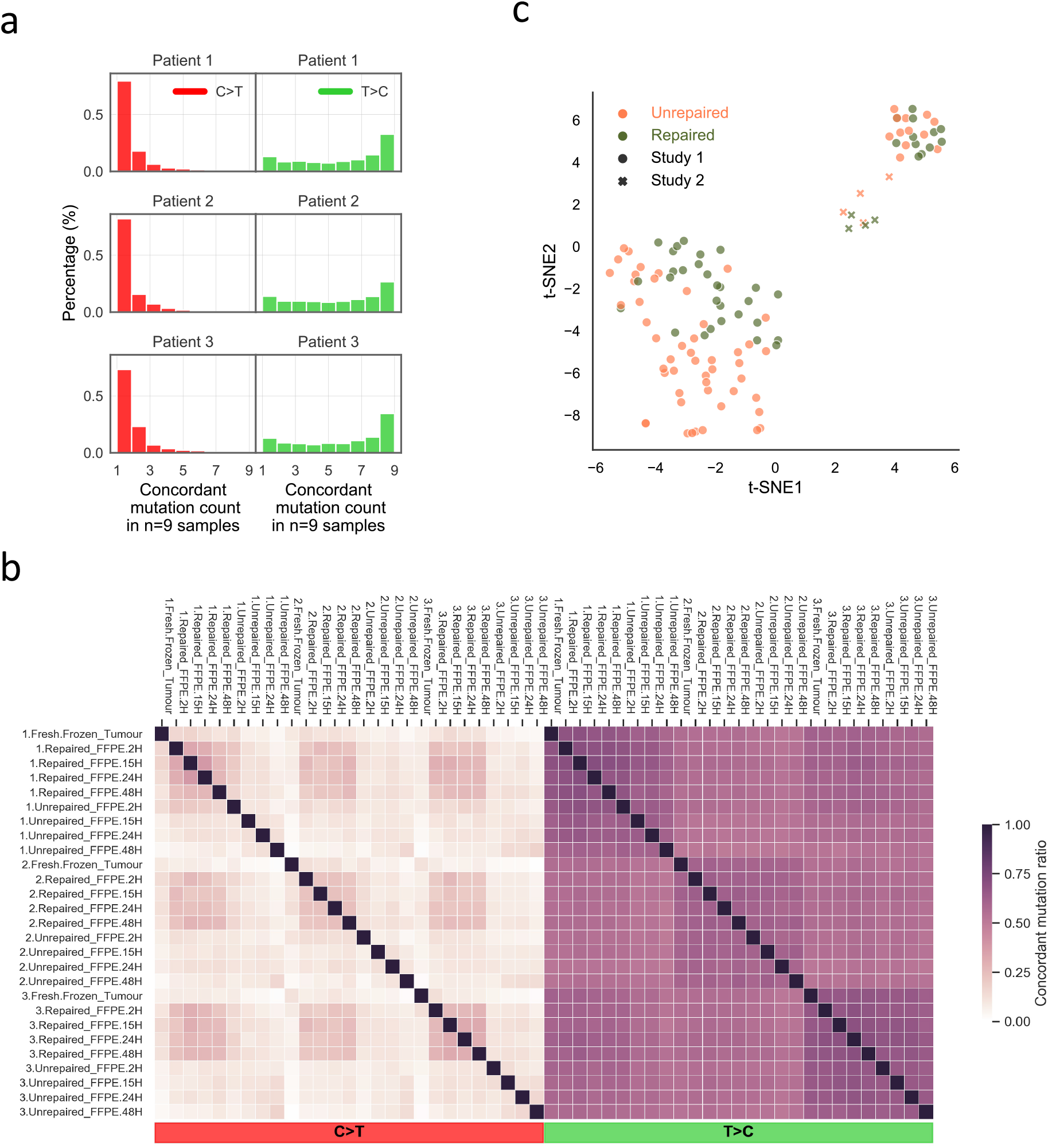
T>C mutations are highly repeated among samples with no specific error profile. We use all mutation data of fixation group (n=27) from study 1 for (a) and (b) as T>C are only over-represented in study 1. We used FFPE-only T>C mutations of all FFPEs (n=110) from study 1 and 2 in (c). (a) Normalised histogram of concordant mutation count per patient. We take all T>C and C>T mutations from the whole mutation list and counted the occurrences for the unique set of all mutations among all samples from each patient (n=9; 4 repaired FFPE + 4 unrepaired FFPE + 1 FF). (b) Pair-wise comparison of concordant mutation ratios for all samples from three patients (n=27). Concordant mutation ratio is calculated using concordant mutation numbers of a sample–pair divided by unique mutation count in the sample pair. (c) Clusters of T>C mutation profiles over 110 FFPE samples. It is the same plot as Fig 1b but using 16-channel of T>C mutation data whereas Fig 1b using 80-channel without T>C mutations. The cluster is represented by t-SNE on cosine metric of 16-channel T>C mutational profiles which are normalized using targeted-region mutational opportunities and whole genome mutational contexts (see Methods & Materials).

**Supplemental Fig4.**
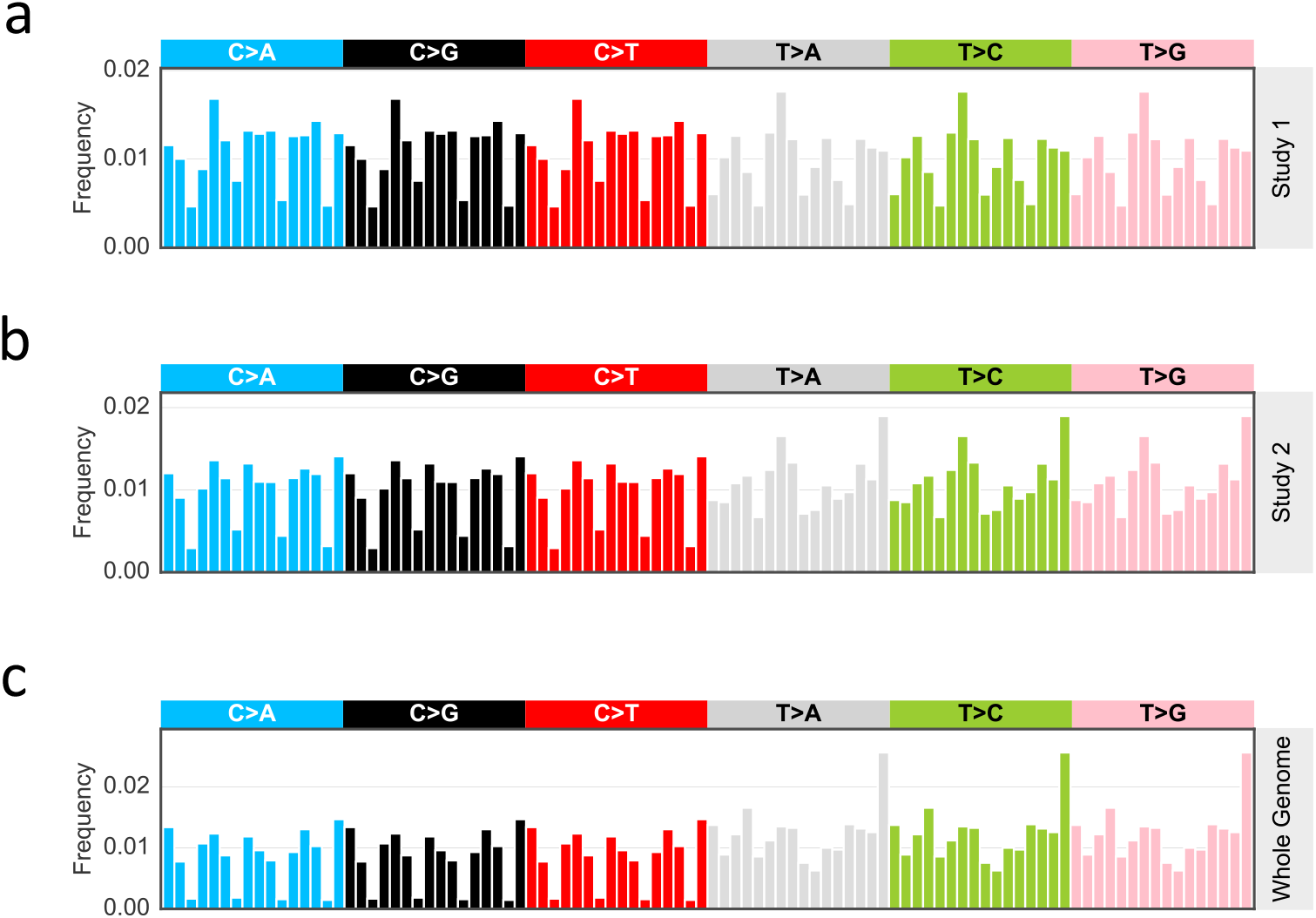
Mutational opportunities. (a) of study 1 targeted regions (b) of study 2 targeted regions (c) of whole genome sequence context

**Supplemental Fig5.**
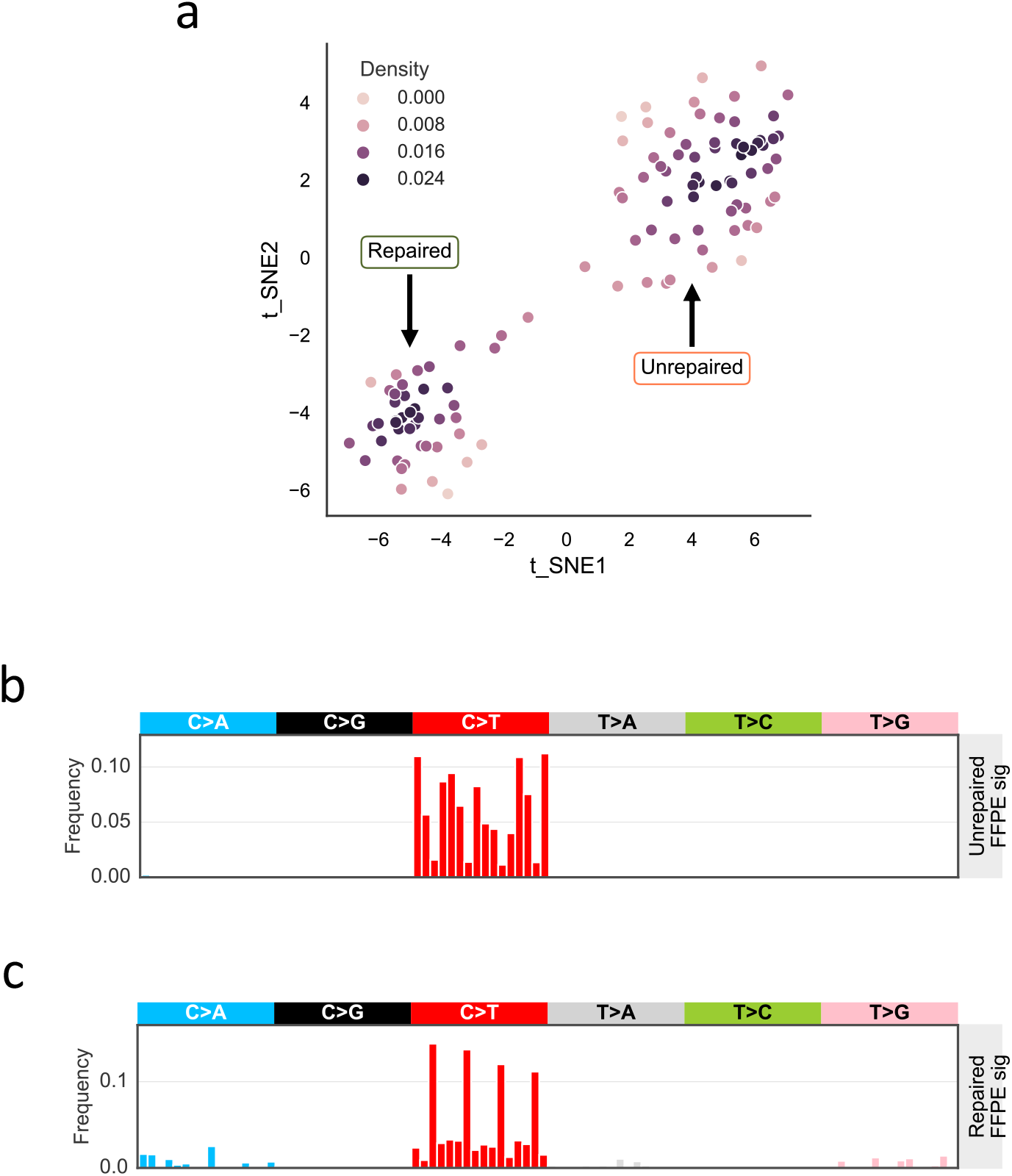
Deriving FFPE signatures from well-representative samples from t-SNE clustering result. (a) Scatter plot of spatial density of t-SNE clustered samples measured using gaussian kernel. The t-SNE cluster is the same as Fig 1b but with spatial density instead. Samples with density value over 0.018 are classified as well-representative samples, and one FFPE signature candidate are generated by averaging the mutational channels. (b) Final version of unrepaired FFPE signature. We repeated (a) for 100 times using different random seeds, thus we have 100 unrepaired FFPE signature candidates. The final version of unrepaired FFPE signature takes the averaged values of all 100 candidates. (c) Final version of repaired FFPE signature. It is derived from the same method as used in (b).

**Supplemental Fig6.**
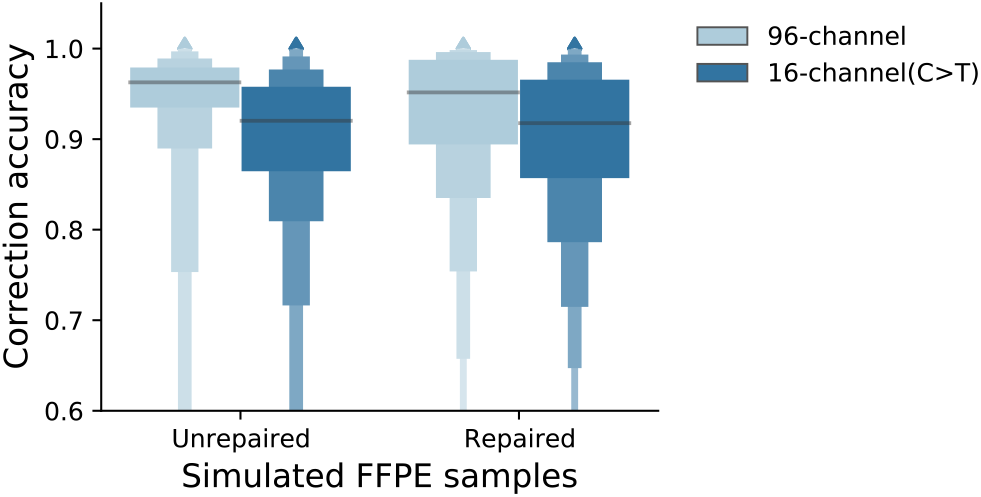
Comparison of correction accuracy measured using all mutations (96-channel) versus using C>T mutations.

**Supplemental Fig7.**
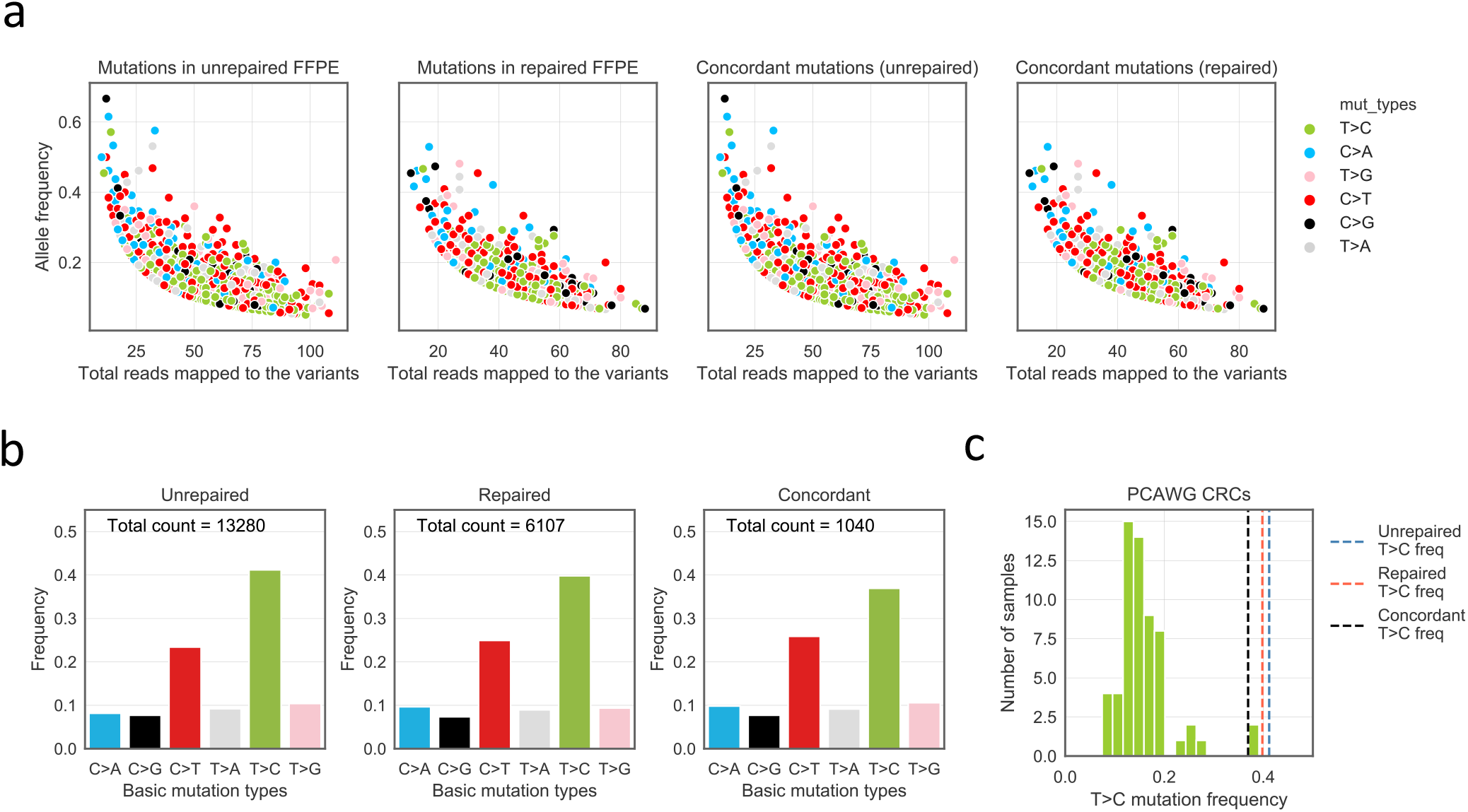
Mutations from two WGS FFPE CRC samples. (a) Allele frequency versus total reads number of detected variants. The four panels from left to right show mutations detected from unrepaired FFPE, repaired FFPE and concordant mutations in unrepaired and concordant mutations in repaired FFPEs, respectively. Concordant mutations refer to variants are detected in both repaired and unrepaired FFPEs with at least 5 supporting reads. (b) Total count of SBS variants in unrepaired, repaired and concordant mutations. (c) T>C mutation frequencies of PCAWG CRC samples. Three dash lines indicate T>C mutation frequencies of unrepaired, repaired and concordant mutations from our sequenced FFPE samples.

**Supplemental Fig8.**
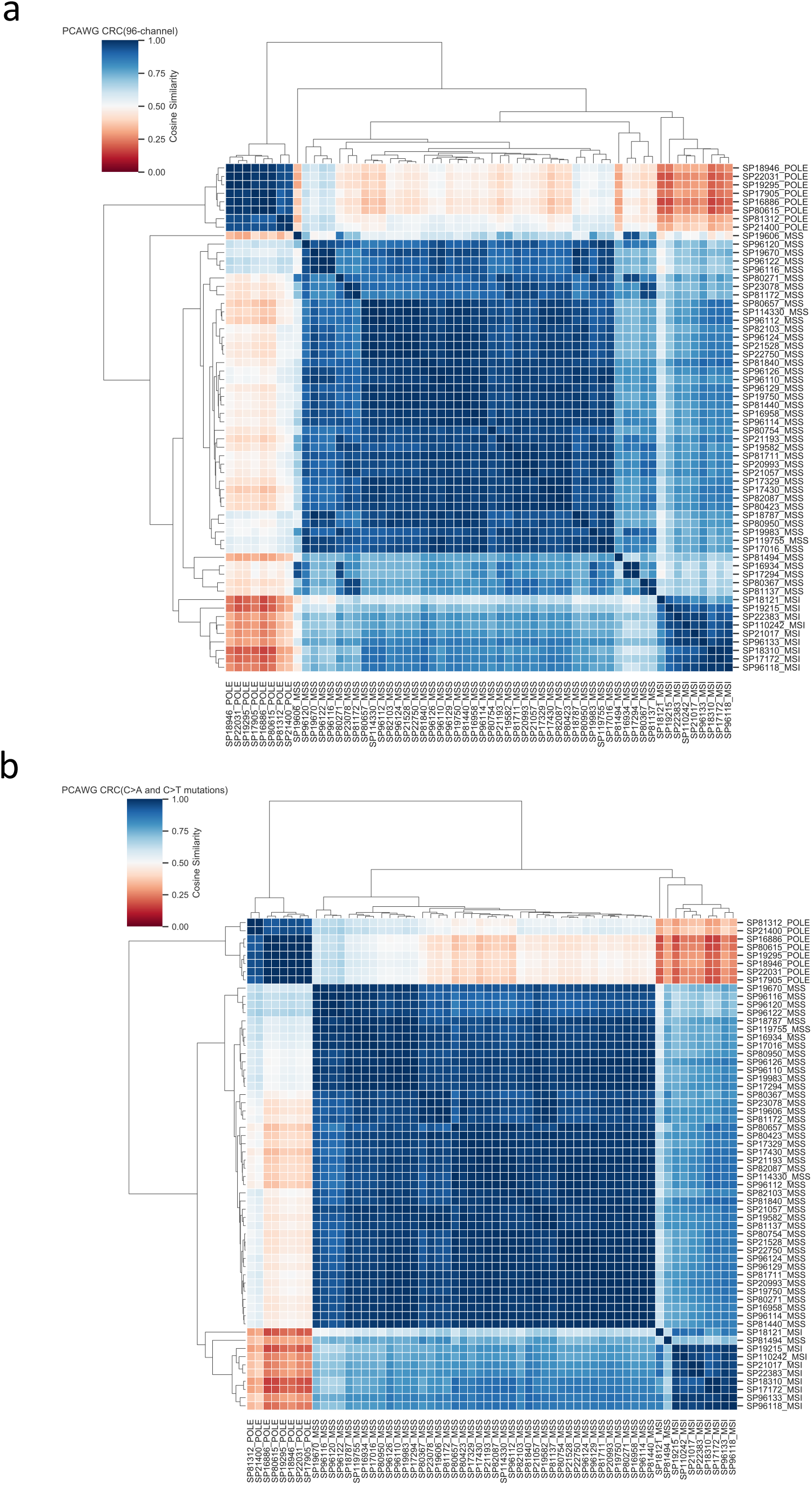
Clustering PCAWG CRC mutational catalogues. (a) using 96-channel profiles. (b) using C>A and C>T mutation profiles.

**Supplemental Fig9.**
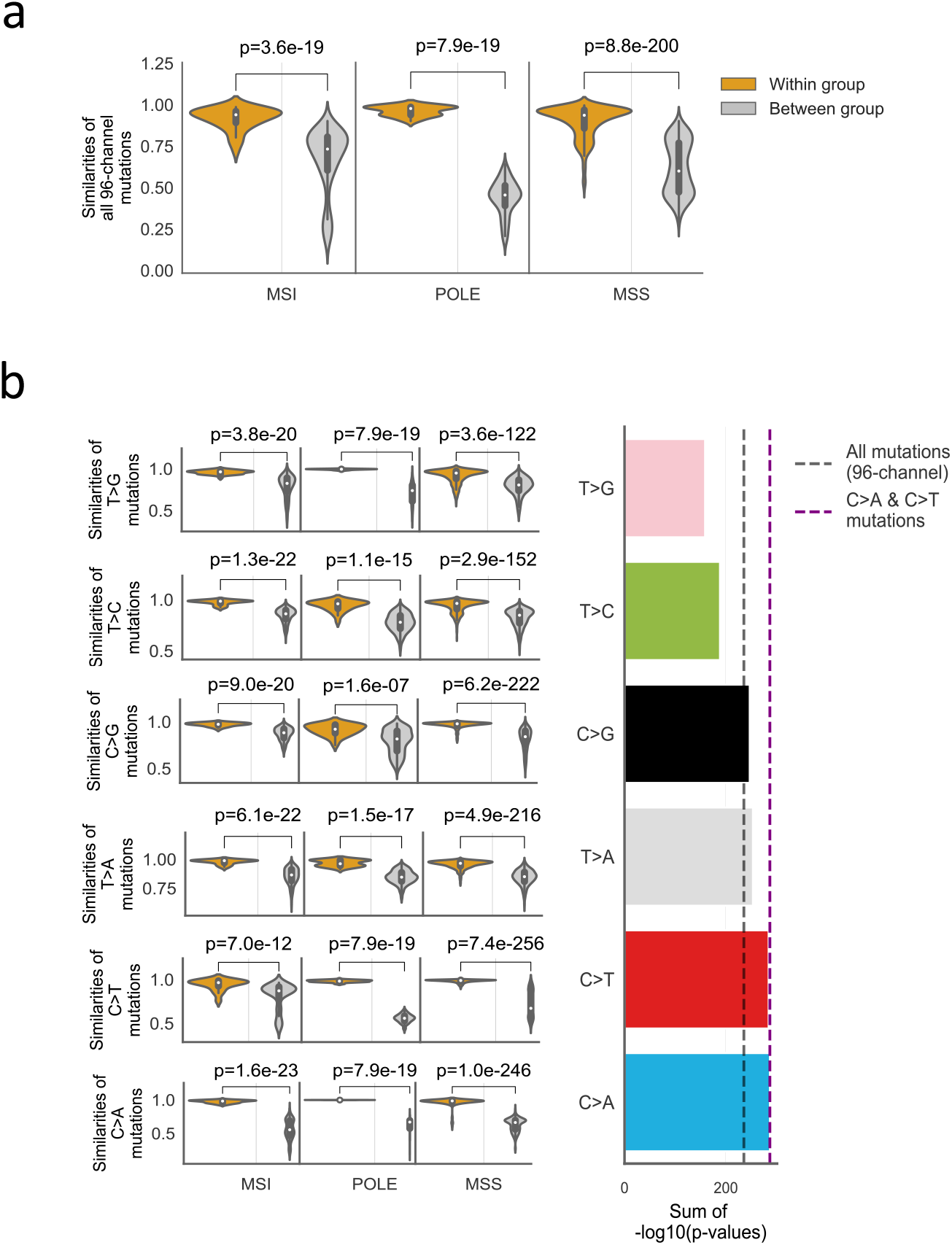
Comparison of sample-pair similarities within and between subgroups of PCAWG CRCs. PCAWG CRC are grouped based on their known labels, namely POLE, MSI and MSS. (a) Comparison made using full 96 channel mutational profiles. The sample-pair cosine similarities of mutation patterns within and between groups are shown in orange and grey box plot, respectively. The difference for each subgroup is measured by two-sided Mann-Whitney *U* test. (b) C>A and C>T mutation patterns are highly conserved/similar within each subtype. The same comparison in (a) is made but using six basic mutation types separately. We use the sum of −log10 (*p*-value) to sort the six mutation types, shown in the right panel. We also use black and purple dash lines to mark sum of −log10 (*p*-value) value by using 96-channel and by using C>A and C>T (32-channel).

**Supplemental Fig10.**
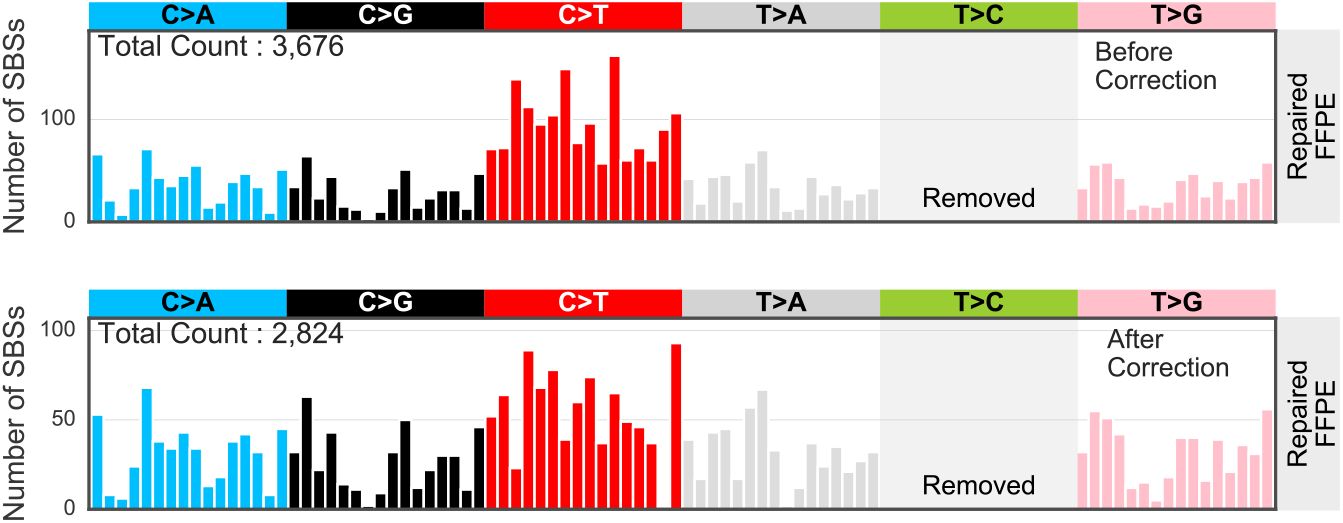
FFPE noise correction results of repaired FFPE CRC sample. The top panel shows mutational profile before correction. And the lower panel shows the corrected profile.

**Supplemental Fig11.**
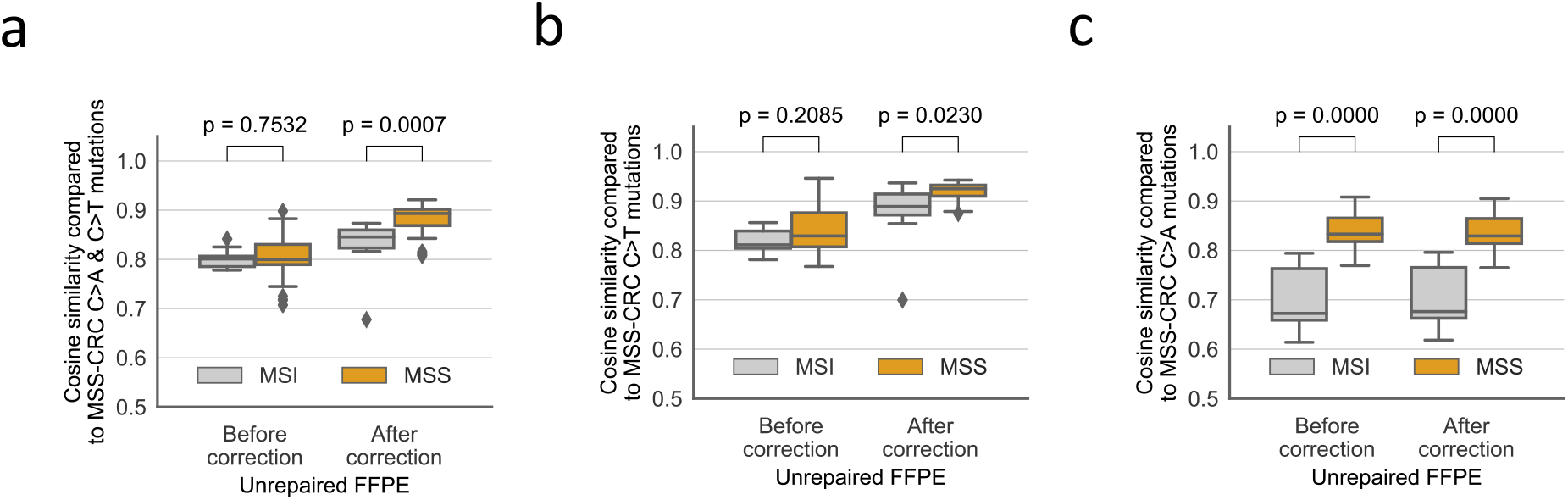
Correction on unrepaired FFPE CRC sample contributes to classify MSS subtype from MSI. The difference for each subgroup is measured by two-sided Mann-Whitney *U* test. (a) Correction makes significant improvement for the classification by using C>A and C>T mutations. (b) Correction on C>T mutations also improves the classification. (c) C>A mutation profiles in unrepaired FFPE sample can also be used as classifier. As our correction acts on C>T channels mostly, so the C>A mutation pattern are almost the same before and after correction (cosine similarity: ~1).

**Supplemental Fig 12.**
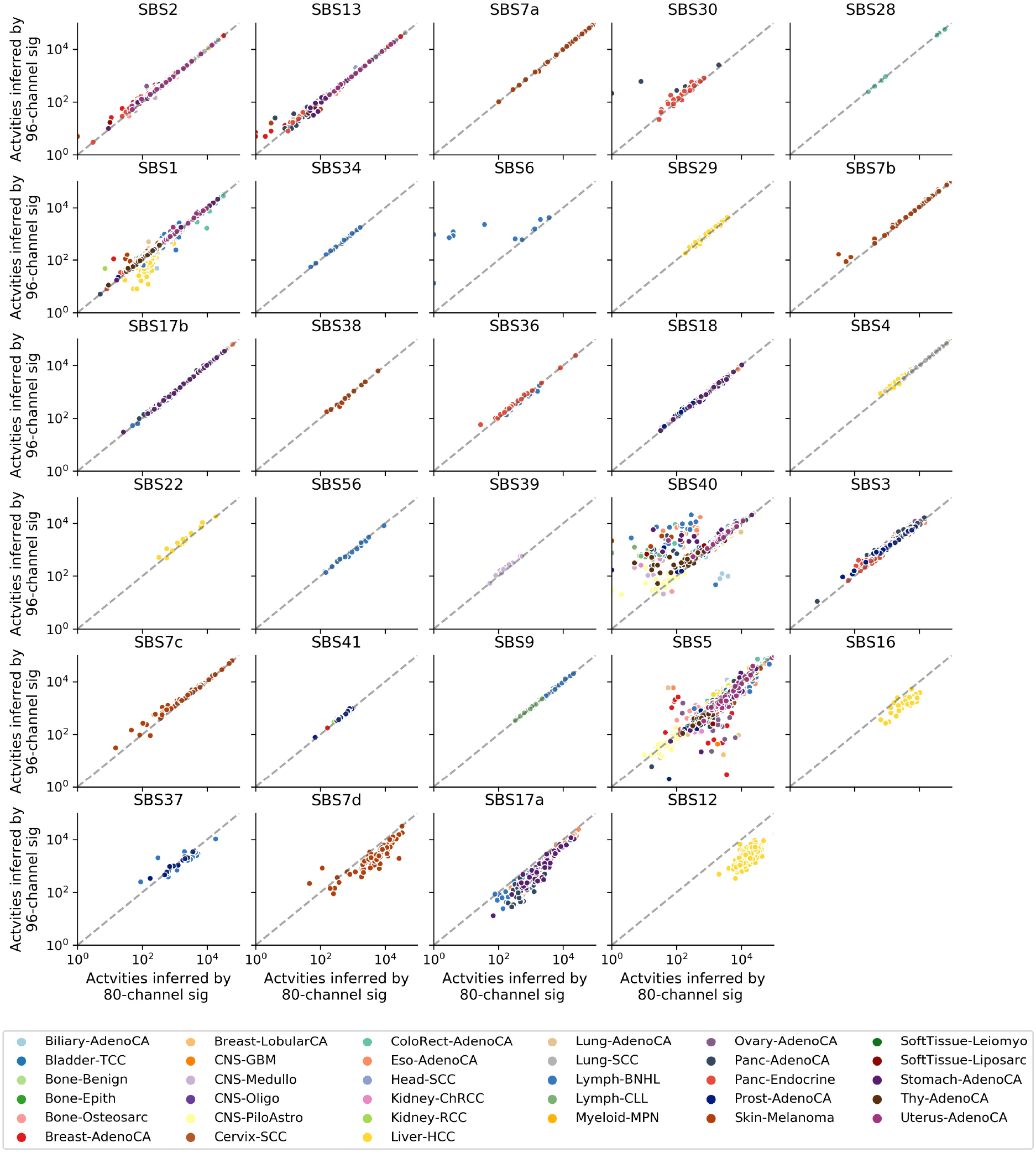
Comparison of refitted activity counts using 80-channel and 96-channel signatures for PCAWG data.

**Supplemental Fig 13.**
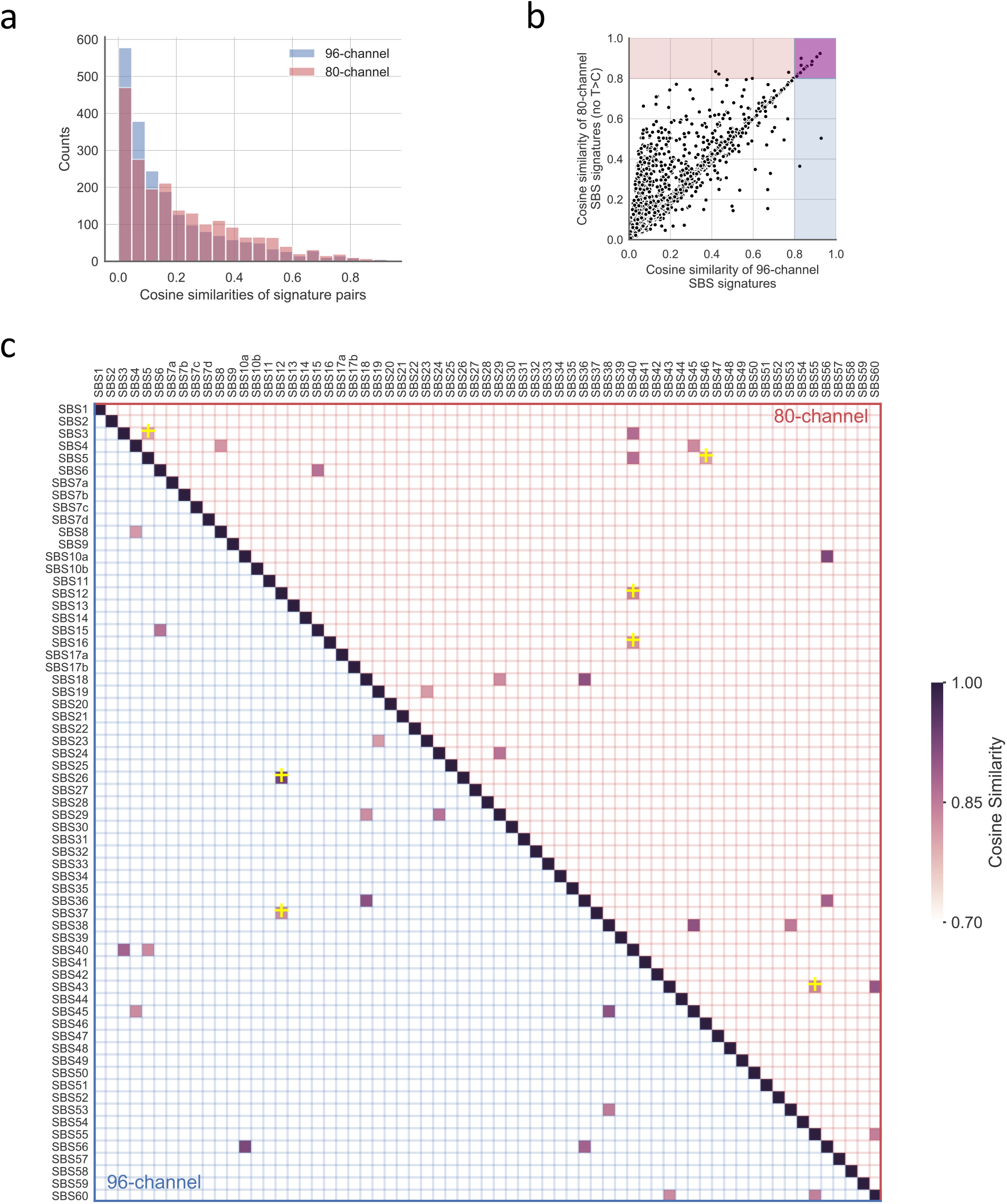
Comparison of signature similarities using 96-channel and 80-channel (no T>C) spectra. (a) Histogram of cosine similarities for signature pairs using 96-channel (96c; blue) and 80-channel (80c; pink). (b) Scatter plot of pair-wise cosine similarities using 96c and 80c signatures. Highly similar (>0.8) signature pairs are highlighted in the plot: 1) purple area shows signature pairs that are highly similar in both signature settings (96c and 80c); 2) blue area contains signature pairs are highly similar by using 96c profiles, but not highly similar by using 80c; and 3) pink area shows pairs with high similarity by using 80c not 96c. (c) Highly similar signature pairs using 96c and/or 80c. The upper and lower triangle show the signature pairs calculated using 80c and 96c, respectively. The signature pair with ‘+’ symbol represents it only exists by using 80c or by using 96c. The pairs with ‘+’ symbol in upper triangle are the dots from pink area in (b), and those in lower triangle are from blue area in (b).

